# Development of blood-cerebrospinal fluid barrier model expressing pharmaceutically important transporters

**DOI:** 10.1101/2024.01.22.576616

**Authors:** Petra Majerova, Krutika Khiratkar, Kevin James, Dominika Olesova, Jozef Vegh, Andrej Kovac

## Abstract

We have established and optimized a protocol for the high-yield isolation of primary epithelial cells from rat choroid plexus. The addition of cytosine arabinoside suppressed the growth of contaminating cells, and epithelial culture was grown into a confluent impermeable monolayer within 5-6 days after seeding. To form an *in vitro* blood-CSF barrier, epithelial cells were plated on inverted coated polycarbonate support of Transwell inserts. Morphologically, the polarized cells remained cuboidal in shape and expressed TJ proteins at a high rate. The filter-grown monolayers displayed transendothelial resistance (TEER) values in the range of 160 to 180 Ω × cm^2^ and remained at this level for 3 days, indicating the persistent formation of continuous TJs. The cells were able to secrete cerebrospinal fluid (CSF) actively. Epithelial cells showed expression of selective influx and efflux transporters. To conclude, our BCSFB model exhibits tight, functional barrier characteristics and shows the functional expression of the pharmaceutically important influx/efflux transporters. The recent model is suitable for *in vitro* investigations of BCSFB and routine pre-clinical drug discovery.

## Introduction

The central nervous system (CNS), as a highly regulated environment, is protected from the periphery by three selective barrier systems: (1) the blood-brain barrier (BBB), (2) the blood-cerebrospinal fluid barrier, (3) the arachnoid membrane. BBB and BCSFB present the main obstacles in the development of pharmacologically active neuropharmaceutics and diagnostics to manage CNS disorders (Pardridge 2005).

The BCSFB is located in brain ventricles and presents an active interface that separates blood from cerebrospinal fluid (CSF) and CSF from the brain (Abbott, Patabendige,. BCSFB is formed by choroid plexus (CP), a highly vascularized structure consisting of single layer of cuboidal epithelial cells, connective tissue and inner layer of endothelial cells that line fenestrated capillaries. Specialized epithelial cells are interconnected by tight junctions (TJs) (Del Bigio 1995). Overall, BCSFB is essential in the regulation of the secretion and composition of CSF, transport of nutrients, and removal of brain metabolites, toxins, and waste products from the CSF. During the inflammation, BCSFB plays important role in the transmigration of peripheral blood cells into the brain through CP epithelium (Blake, Carrigan et al. 2004, Ryan, Grimes et al. 2005, Tenenbaum, Steinmann et al. 2013, Lauer, Marz et al. 2019).

CP has been studied in relation to different pathological conditions including neurodegenerative disorders such as Alzheimeŕs disease (AD), and is also considered as a target for the CNS delivery of therapeutics. Various *in vitro* model platforms have been set up to study the function, phenotype and molecular properties of the BCSFB to simulate disease process and evaluate drug permeability into the brain (Dabbagh, Schroten et al. 2022). Such models mimic the *in vivo* conditions and demonstrate representative properties of TJs, restricted paracellular permeability and expression of specific transporters and receptors (Gaillard, Voorwinden et al. 2001, Schroten, Hanisch et al. 2012, Di Marco, Gonzalez Paz et al. 2019). The *in vitro* models are established in varied range of technical and physiological complexities. Routinely, 2D systems based on the cultivation of choroid plexus epithelial cell monocultures on semipermeable membranes are used for permeability studies, modeling of immune function, evaluation of the toxicological effect of investigated molecules, and structure-activity assays (Shi and Zheng 2005, Lazarevic and Engelhardt 2016). In these models, epithelial cells are usually grown as tight monolayers on culture filter inserts to develop an active cellular barrier. Epithelial cells can be cultivated in standard or inverted filter systems. In the standard *in vitro* model, the cells grown on the upper face of the filter membrane. On the contrary, in the inverted model the cells are cultivated on the lower support of the membrane (Dinner, Borkowski et al. 2016, Lauer, Marz et al. 2019).

At present, BCSFB *in vitro* models based on immortalized cell lines or primary cell cultures are available. The rat CP epithelial cell lines Z310 and TR-CSFB3 have been developed using transfection with the simian virus 40 large T-antigen (Kitazawa, Hosoya et al. 2001, Zheng and Zhao 2002). The human cell line CPC-2 was derived from CP carcinoma (Kumabe, Tominaga et al. 1996). Regarding to primary cultures, CP epithelial cells have been established from small animals, including mice (Thomas, Stadler et al. 1992, Barkho and Monuki 2015), and rat (Villalobos, Parmelee et al. 1997, Zheng, Zhao et al. 1998, Zheng and Zhao 2002) and large animals such as rabbit (Ramanathan, Hui et al. 1996), pig (Hakvoort, Haselbach et al. 1998, Angelow, Zeni et al. 2004, Baehr, Reichel et al. 2006), cow (Crook, Kasagami et al. 1981), sheep (Holm, Hansen et al. 1994), canine (Hu, Bian et al. 2014, Braun, Sakamoto et al. 2017), and non-human primates (Delery and MacLean 2019). Human primary CP epithelial cultures have been obtained from embryos, or postmortems (Ishiwata, Ishiwata et al. 2005, Niehof and Borlak 2009). Compared to immortalized cell lines that are more accessible and easy to culture, primary cultures are usually expensive and laborious to isolate but reliably mimic the *in vivo* state.

The development of *in vitro* systems that can modulate real *in vivo* conditions is necessary and critical for further neuroscience research. Therefore, novel and advanced model systems should be established. Here, applying the optimized isolation and cultivation protocol, we established a primary rat inverted *in vitro* model of BCSFB. We analyzed the expression of epithelial cell markers, maturation of tight junctions, and functional barrier characteristics. We described the expression of a pharmaceutically important influx/efflux transporter to demonstrate the suitability of our model for pre-clinical drug discovery.

## Material and methods

### Animals

All animals used in this study were from the in-house breeding colony (SPF like, monitored according to the Federation of European Laboratory Animal Science Associations). In this study, we used Sprague Dawley rats. Animals were housed under standard laboratory conditions with free access to water and food and were kept under diurnal lighting conditions. All experiments on animals were carried out according to the institutional animal care guidelines conforming to international standards (Directive 2010/63/EU) and were approved by the State Veterinary and Food Committee of Slovak Republic (RO-1101/14-221C). All animal experiments were monitored by the Institutional Ethics Committee (Ethics Committee of the Institute of Neuroimmunology).

### Isolation of choroid plexus epithelial culture from rat (CPEC-R)

The epithelial culture was prepared from the choroid plexus of 7-day-old Sprague Dawley rats (n = 12/isolation). The whole brains were placed in ice-cold DMEM-F12 medium (PAA Laboratories GmbH, Germany) with gentamycin (Sigma-Aldrich, St. Louis, MO) to chill the tissue and wash off the blood. The choroid plexuses were dissected from both lateral and third ventricles. The tissue was centrifuged for 10 min at 800xg. For digestion, we used the Pronase enzyme (Roche Diagnostics, Indianapolis, USA) at the final concentration of 1.5 mg. mL^-1^ enzyme in DMEM/F12 medium and incubated for 10 min at 37°C. The tissue was incubated with a pre-prepared digestion mix at 37°C for 10 min with gentle shaking. The reaction was stopped when 10 mL of complete medium was added to the digestion mixture. The tissue was centrifuged for 10 min at 800xg. After the centrifugation, the cells were mechanically dissociated by passages through a 20-gauge needle in 5 mL of complete media. An aliquot of cell suspension was removed and mixed with 20 μL of 0.4% Trypan blue (Sigma-Aldrich, St. Louis, MO) to count cell number and to assess the viability. The cells were cultivated in DMEM-F12 medium containing 20% fetal bovine serum (FCS, Thermo Fisher Scientific, Waltham, Massachusetts, USA), 2 mM L-glutamine (Life Technologies Invitrogen, Carlsbad, CA), BME vitamins (Sigma-Aldrich, St. Louis, MO), Insulin-Transferrin-Selenium (Life Technologies Invitrogen, Carlsbad, CA), 3 µM epidermal growth factor (EGF, Sigma-Aldrich, St. Louis, MO), 0.1% hydrocostisone (Sigma-Aldrich, St. Louis, MO), 20 mM cytosine β-D-arabinofuranoside (Ara-C, Sigma-Aldrich, St. Louis, MO) and gentamycin at 37°C, 5% CO_2_ in a water-saturated atmosphere.

### Development of inverted primary rat model of BCSFB

Inserts were pre-coated with 10 μg/cm^2^ collagen type IV (Sigma-Aldrich, St. Louis, MO) and 5 μg/cm^2^ fibronectin (Sigma-Aldrich, St. Louis, MO). 200 μL of cell suspension was plated onto the top of the basolateral site of 0.4 μm polycarbonate Transwell inserts (Becton Dickinson, New Jersey, USA). After 24 hours the inserts were turned over and placed in a plate with complete medium. The cells were cultivated in DMEM-F12 medium containing 10% FCS, 2 mM L-glutamine, BME vitamins, Insulin-Transferrin-Selenium and 0.1% hydrocortisone at 37°C, 5% CO_2_ in a water-saturated atmosphere. The cells were cultivated for 5-6 days. The medium was changed every 2 days.

### Transepithelial electrical resistance (TEER) and epithelial permeability

TEER values were determined, using the STX-2 electrode system (World Precision Instruments, Berlin, FRG). The final TEER values were multiplied by the filter surface area. To determine the epithelial permeability of Lucifer Yellow (LY), Transwell inserts (in a 12-well format, containing an epithelial monolayer or without cells) were transferred into 12-well Costar plates containing 1.5 mL of Ringer-HEPES solution (150 mM NaCl, 5.2 mM KCl, 2.2 mM CaCl_2_, 0.2 mM MgCl_2_, 6 mM NaHCO_3_, 5 mM HEPES, 2.8 mM glucose; pH 7.4) per well. The cell culture medium was removed from the inserts, and 0.5 mL of Ringer-HEPES solution containing 10 μM LY (Sigma-Aldrich, St. Louise, MO) was added to the lower (abluminal) compartment. All incubations were performed at 37°C. After 5, 10, 15, 30, and 60 min, 200 μL aliquots from each upper compartment was quantified for fluorescence intensity (Fluoroscan Ascent FL, Labsystems; excitation wavelength: 428 nm; emission wavelength: 536 nm). Control values were obtained from filters without cells. The permeability coefficient (Pe) of LY was calculated in cm/s. The permeability values of the inserts (PSf, for inserts with a coating only) and the inserts plus epithelium (PSt, for inserts with a coating and cells) were taken into consideration by applying the following equation: 1/PSe = 1/PSt − 1/PSf. To obtain the epithelial permeability coefficient (Pe, in cm/s), the permeability value corresponding to the epithelium alone was then divided by the insert’s porous membrane surface area.

### Measurements of CSF secretion

Cells were washed in CSF secretion buffer (122 mM NaCl, 4 mM KCl, 1 mM CaCl_2_, 1 mM MgCl_2_, 15 mM NaHCO_3_, 15 mM HEPES, 0.5 mM Na_2_HPO_4_, 0.5 mM NaH_2_PO_4_, 17.5 mM glucose, 5 µg. mL^-1^ insulin, pH 7.3)(Hakvoort, Haselbach et al. 1998). Before the experiment, the cells were pre-incubated for 1hour with secretion buffer. Subsequently, 500 µl or 1.5 mL of 1µM 70 kDa fluorescent Texas Red dextran (70 kDa dextran impermeable to the cell monolayer, Sigma-Aldrich, St. Louis, MO) was added to apical and basolateral compartments. The secreted CSF volume (µL/cm^2^) was calculated as a function of dextran concentration change. Filters were incubated at 37°C up to 16 hours. 200 µL samples were taken from the apical and basolateral chamber and replaced with fresh dextran solution. Samples were analysed in a fluorescent plate reader (Fluoroskan Ascent, Thermo Fisher Scientific, Pittsburgh, United States). CSF volumes secreted were corrected and calculated according to the equation (Hakvoort, Haselbach et al. 1998):

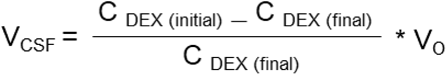

Where, VCSF = CSF volume (μl), CDEX (initial) = initial (μM), CDEX (final) = final (μM), V0 = initial volume applied (μl).

### Immunocytochemical staining

CPEC-R were fixed for 10 min in ice-cold acetone/ethanol solution (8:2), wash with phosphate buffer saline (PBS, 137 mM NaCl, 2.7 mM KCl, 10 mM Na_2_HPO_4_, 2 mM KH_2_PO_4_, pH 7.4) and blocked for 60 min in 5% bovine serum albumin (BSA, Sigma-Aldrich, St. Louis, Missouri, United States) in PBS. Sections were incubated in primary antibodies (see Table 1). After washing, the sections were incubated 1 hour in secondary antibodies: goat anti-rabbit or goat anti-mouse AlexaFluor488/546 (1:1000; Invitrogen Life Technologies, Carlsbad, CA). Sections were mounted (Sigma-Aldrich, St. Louis, Missouri, United States) and examined using a LSM 710 confocal microscope (Zeiss, Jena, Germany).

**Table 1:** List of used primary antibodies.

| Primary antibody | Reactivity | Dillution (WB) | Dillution (ICH) | Source |
| --- | --- | --- | --- | --- |
| Cytokeratine | monoclonal mouse anti-rat | 1:1000 | 1:1000 | Abcam |
| Claudin-2 | polyclonal rabbit anti-rat | 1:500 | 1:250 | Abcam |
| Claudin-1 | polyclonal rabbit anti-rat | 1:500 | 1:250 | Abcam |
| Occludin | polyclonal rabbit anti-rat | 1:500 | 1:250 | Thermo Scientific |
| ZO-1 | polyclonal rabbit anti-rat | 1:1000 | 1:1000 | Thermo Scientific |
| LRP-1 | polyclonal rabbit anti-rat | 1:500 | 1:500 | Thermo Scientific |
| LRP-2 | polyclonal rabbit anti-rat | 1:500 | 1:500 | Thermo Scientific |
| Pgp-1 | polyclonal rabbit anti-rat | 1:500 | 1:500 | Thermo Scientific |
| LAT-1 | polyclonal rabbit anti-rat | 1:500 | 1:500 | Thermo Scientific |
| BRCP1 | polyclonal rabbit anti-rat | 1:500 | 1:500 | Thermo Scientific |
| MRP2 | polyclonal rabbit anti-rat | 1:500 | 1:500 | Thermo Scientific |
| MRP3 | polyclonal rabbit anti-rat | 1:500 | 1:500 | Thermo Scientific |
| MRP5 | polyclonal rabbit anti-rat | 1:500 | 1:500 | Thermo Scientific |
| RAGE | polyclonal rabbit anti-rat | 1:1000 | 1:500 | Abcam |

### Western blot analysis

Epithelial cells and choroid plexus tissue were harvested into lysis buffer (200 mM Tris, pH 7.4, 150 mM NaCl, 1 mM EDTA, 1 mM Na_3_VO_4_, 20 mM NaF, 0.5% Triton X-100, 1 × protease inhibitors complete EDTA-free from Roche, Mannheim, Germany). Total protein concentration of prepared cell extracts was measured by Bio-Rad protein assay (Bio-Rad laboratories GmbH, Colbe, Germany). A total of 30 μg of the proteins was separated onto 10% SDS-polyacrylamide gels and transferred to nitrocellulose membrane in 10 mM N-cyclohexyl-3-aminopropanesulfonic acid (CAPS, pH 11, Roth, Karlsruhe, Germany). The membranes were blocked in 5% milk in Tris-buffered saline (TBS-T, 137 mM NaCl, 20 mM Tris-base, pH 7.4, 0.1% Tween 20) for 1 hour and incubated with primary antibody (Table 1) overnight at 4°C. Membranes were incubated with horseradish peroxidase (HRP)-conjugated secondary antibody in TBS-T (1:3000, Dako, Glostrup, Denmark) for 1 hour at room temperature. Immunoreactive proteins were detected by chemiluminescence (SuperSignal West Pico Chemiluminescent Substrate, Thermo Scientific, Pittsburgh, United States) and the signals were digitized by Image Reader LAS-3000 (FUJIFILM, Bratislava, Slovakia). The signal was semiquantified by ImageJ.

### Mass spectrometry

200 µl of secretome was precipitated by acetone overnight at -20 °C. The samples were centrifuged and pellet dissolved in 8 M urea in Tris-HCl, pH= 8. The total protein concentration of prepared tissue extracts was measured by Bio-Rad protein assay (Bio-Rad Laboratories GmbH, Colbe, Germany). 100 µg of proteins were reduced with 10 mM dithiothreitol (Sigma-Aldrich, MO, USA) at 37 °C for 60 min. Alkylation was performed with 100 µL of 50 mM iodoacetamide (Sigma-Aldrich, MO, USA) in the dark for 30 min. Proteins were digested with trypsin (Promega, Wisconsin, USA) in a ratio of 1:100 at 37 °C, overnight. Aliquots of purified complex peptide mixtures of 100 ng were separated using Acquity M-Class UHPLC (Waters, Milford, MA, USA). Samples were loaded onto the nanoEase Symmetry C18 trap column (25 mm length, 180 μm diameter, 5 μm particle size). After 2 min of desalting/concentration by 1% acetonitrile containing 0.1% formic acid at a flow rate of 8 μL/min, peptides were introduced to the nanoEase HSS T3 C18 analytical column (100 mm length, 75 μm diameter, 1.8 μm particle size). For the thorough separation, a 90 min gradient of 5%–35% acetonitrile with 0.1% formic acid was applied at a flow rate of 300 nL/min. The samples were sprayed (3.1 kV capillary voltage) to the quadrupole time-of-flight mass spectrometer Synapt G2-Si (Waters, Milford, MA, USA). Spectra were recorded in a data-independent manner in high-definition MSe mode. Ions with 50–2000 m/z were detected in both channels, with a 1 s spectral acquisition scan rate. Data processing was done in Progenesis QI 4.0 (Waters). For peak picking, the following thresholds were applied: low energy 320 counts and high energy 40 counts. Precursors and fragment ions were coupled, using correlations of chromatographic elution profiles in low/high energy traces. Then, peak retention times were aligned across all chromatograms. Peak intensities were normalized to the median distribution of all ions, assuming the majority of signals are unaffected by experimental conditions.

### ImageJ

ImageJ was used for the evaluation and quantification of chemiluminescence and immunocytochemistry slides. Relative staining patterns and intensity of projections were visualized by confocal microscopy and evaluated by image analysis. We quantified 10 slices from each sample. For semiquantitative analysis, the color pictures were converted to grayscale 8-bit TIFF file format, and regions of interest were analyzed with ImageJ software. The grayscale 8-bit images were converted to thresholded 1-bit images, on which the number of immunolabeled structures localized in the area of interest was measured. Then the average intensity/pixel values of each area were calculated, and the average intensity/pixel values representing the background intensity were subtracted from those of immunolabeled areas.

### Data analysis and statistics

Each experiment was repeated at least three times. Data are presented as mean ± standard deviation (SD). Differences between means were analyzed with an independent two-tailed Student’s t-test and one-way ANOVA (Prism 8.0 software, GraphPad, Inc., SanDiego, CA). Differences at p<0.05 were regarded as statistically significant. Differential expression (DE) analysis was performed using the R software environment (version 4.3.1) (https://www.r-project.org). Data were further transformed using natural logarithm (ln), and DE proteins (p < 0.01) were identified using a pairwise two-sided t-test assuming unequal variances (Welch test). Top 50 differentially expressed proteins (ANOVA, p>0.01) are shown in heatmap, which was generated using the pheatmap package in R. Gene Ontology (GO) enrichment analysis of DE proteins (ANOVA, p>0.01) was performed in R using packages clusterprofiler, and enrichplot (Wu, Hu et al. 2021).

## RESULTS

### Functional barrier characteristics of BCSFB model formed by CPEC-R

Epithelial cells were dissociated from choroidal tissue by pronase digestion and cultured in standard DMEM media supplemented with 20% FCS. We showed that after pronase treatment, the cell viability was 84 ± 5%. For comparison, lower viability was demonstrated after collagene/dispase (74± 5.8%) and trypsine digestion (61± 7.4%). After pronase treatment, we obtained 3–5 × 10^5^ cells from the choroid plexuses of 12 rats (Figure 1A). Following isolation, epithelial cells were plated on coated polycarbonate lower support of Transwell inserts (pore size 0.4 µm, surface area 1.12 cm) at seeding density 1. 10^4^ cells per mL (Figure 1B). The addition of cytosine arabinoside suppressed the growth of contaminating cells, and the epithelial culture developed into a confluent monolayer within 5-6 days. To visualize actin distribution, we stained the culture with fluorescent-labelled phalloidin (FITC-phalloidin), which binds specifically to f-actin. Actin filaments are associated with TJs and are directly involved in controlling paracellular permeability (Chifflet and Hernandez 2012). DAPI was used for cell nuclei staining. Staining revealed actin filaments in networks and bundles. Figure 1C showed that cells in collagen remained cuboidal in shape. Further we characterized and quantified epithelial barrier function by two standard methods: TEER and paracellular permeability to LY. Before measuring TEER, we removed the cell cultures from the incubator and equilibrated them to room temperature. This allowed higher consistency in TEER values, which reached a maximum of 5-6 days in culture. The average TEER values were 172 ± 9.5 Ω × cm^2^ and remained at this level for 3 days, indicating the persistent formation of continuous TJs (Figure 1D). We also showed that steroids (dexamethasone, hydrocortisone) modulated the TEER values. We stimulated epithelial monolayer using 1 µM dexamethasone or 2.8 mM hydrocortisone and observed an increase in TEER. Dexamethasone increased TEER from 185.2 ± 12.26 to 226.7 ±13.5 Ω × cm^2^ (n=6, p= 0.0005). Hydrocortisone increased TEER from 185.2 ± 12.26 to 217.3 ± 21 Ω × cm^2^ (n=6, p=0.015). On the other hand, change to serum-free medium did not increase or lowered the epithelial resistance (159.8 ± 32.34 Ω × cm^2^, n=6, p= 0.13, Figure 1E).

**Figure 1.**
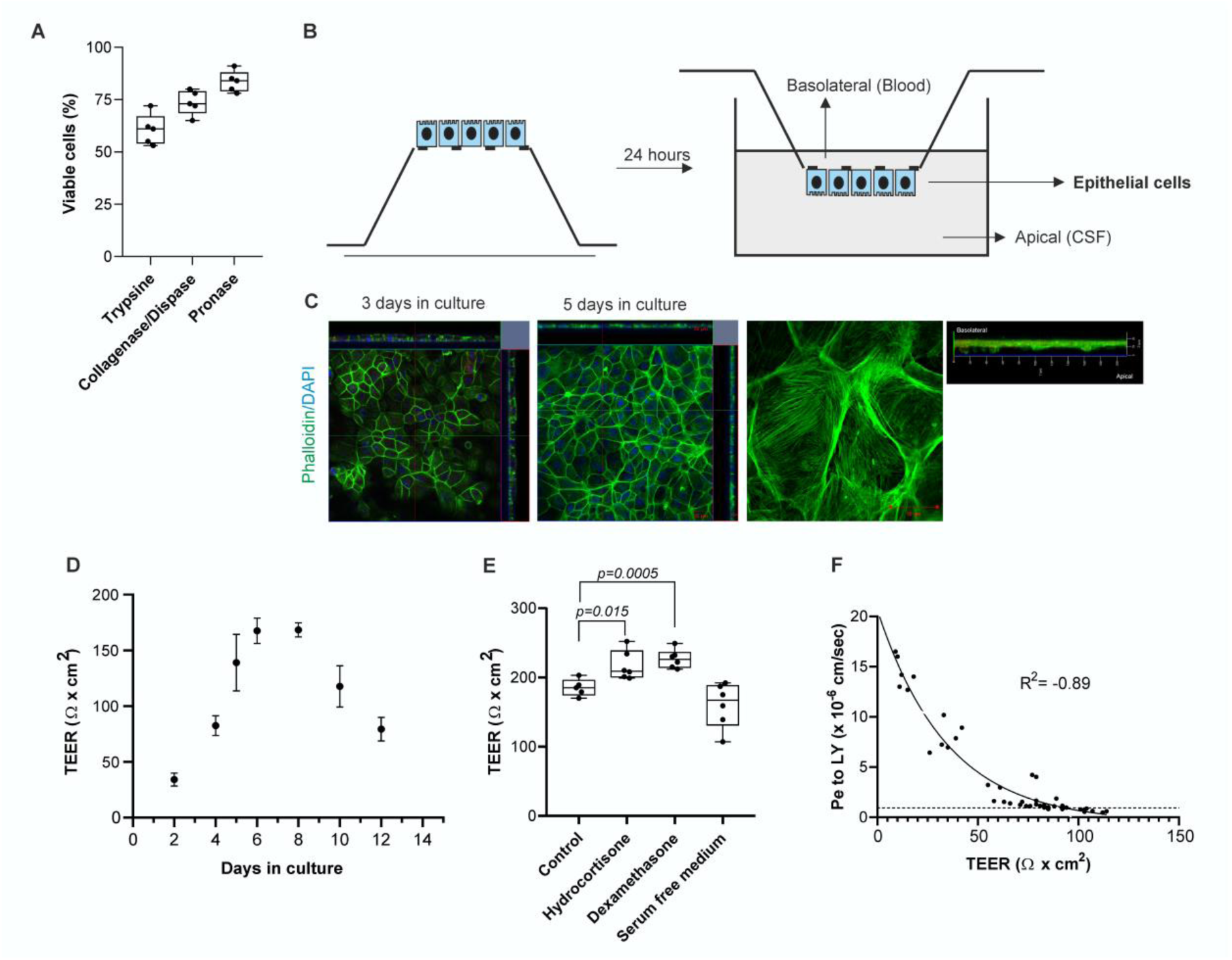
Establishment of inverted *in vitro* BCSFB model. (**A**) Cell viability after enzymatic treatment (the data are presented as mean ± SD). (**B**) The BCSFB model was formed by CPEC-R seeded on the inverted side of Transwell inserts. (**C**) The cuboidal morphology of epithelial cells was imaged by fluorescence microscopy. The actin of epithelial cells was stained with Alexa Fluor 488 phalloidin. The labelled epithelial monolayer was scanned by confocal microscopy to show the side view (x-z plane) of the barrier. Scale bar=20 and 50 µm. (**D**) TEER was measured following equilibration of the Transwell inserts to room temperature. TEER was expressed as Ω × cm^2^ (the data are presented as mean ± SD, n=5). (**E**) Dexamethasone and hydrocortisone stimulated increase in TEER (the data are presented as mean ± SD, n=5). P < 0.05 was considered to be statistically significant. * p < 0.05, ** p ≤ 0.01 and *** p ≤ 0.001. (**F**) Permeability of LY through epithelial monolayer. Pe was calculated over a 90 minutes time-course at 37°C. Correlation between TEER and permeability to LY over 90 minutes. Data was fitted to a one-phase exponential decay curve (the data are presented as mean ± SD, n=5).

Next, we quantified paracellular permeability to LY over 90 minutes. The LY permeability coefficient was 0.95 ± 0.13 × 10^-6^ cm/sec^-1^. Furthermore, a strong correlative relationship was observed between the TEER values and permeability to LY. The relationship fitted an exponential decay curve (R^2^ = -0.89), showing that as TEER increased, Pe to LY significantly decreased (Figure 1F).

### CPEC-R actively secreted CSF

To measure CSF secretion, Texas Red dextran was added to the abluminal compartment of the BCSFB model, and secreted CSF volume (in µL/cm^2^) was calculated as a function of dextran concentration change. Figure 2 shows the CSF volume secreted hourly by cultured epithelial cells for up to 14 hours. The volume of CSF secreted by the monolayer increased in linear fashion, reaching a plateau after 12 hours of approximately 136 ± 29.2 µL/cm^2^. No increase in CSF volume secreted per time interval was seen after 12 hours. Using measurements made during the first 4 hours of linear CSF secretion, CSF production rates were calculated to be 21.8±8.1 μl/cm^2^ /h.

**Figure 2.**
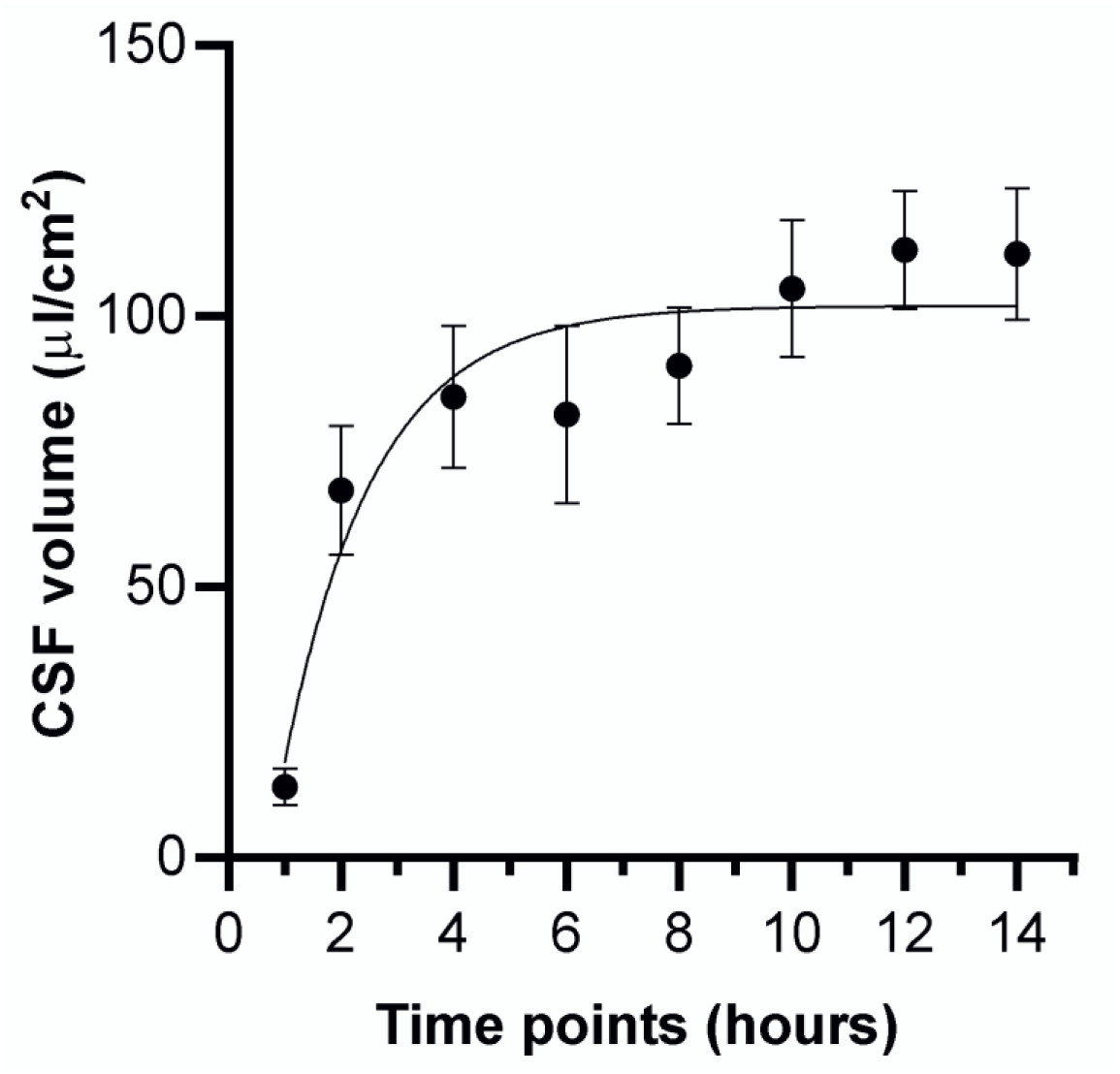
CPEC-R secreted CSF. CSF secretion rates were calculated from the dilution of Texas Red dextran. Measurements were taken every 2 hours up to 14 hours. The presented values are mean ± SD (n=5).

### Inverted *in vitro* BCSFB barrier developed morphologically dense tight junctions

TJ proteins are crucial for cellular polarization and regulation of barrier function. TJs are present at the apical side of epithelium and play an important role in the establishment and maintenance of barrier function. To investigate the presence of TJ components we performed immunocytochemical and western blot analysis of claudin-1, claudin-2, occludin and ZO-1 expression. To gain insight regarding the tight junction morphology of CPEC-R we analyzed cell layers grown inverted on Transwell inzerts by immunofluorescence staining (Figure 3A). Fluorescent images of cultured cells revealed the monolayer structure of the confluent epithelial culture. All investigated TJs proteins displayed intense staining and a continuous pattern localized apically at the sites of intercellular contact, suggesting the formation of highly organized, continuous TJs. The epithelial marker-pan cytokeratine was localized throughout the cytoplasm. Constitutive expression of pan cytokeratin showed that the cells are naturally committed toward an epithelial phenotype. All investigated proteins were also detected on protein level (Figure 3B). Western blot analysis demonstrated, that the expression levels of ZO-1 was comparable to whole CP tissue (CP: 4521 ± 1905, CPEC-R: 6733 ± 1015, p= 0.36). Expression of claudin-1 (CP: 18617 ± 494, CPEC-R: 11638 ± 609.2, p= 0.009) and occludin (CPEC-R: 3789 ± 158, CP: 3111 ± 149.6, p= 0.035) was slightly decreased in cell culture. On the other hand, claudin-2 expression was significantly lower in the *in vitro* culture (CP: 8904 ± 813.9, CPEC-R: 3964.9 ± 688.6, p= 0.0007, Figure 3C)

**Figure 3.**
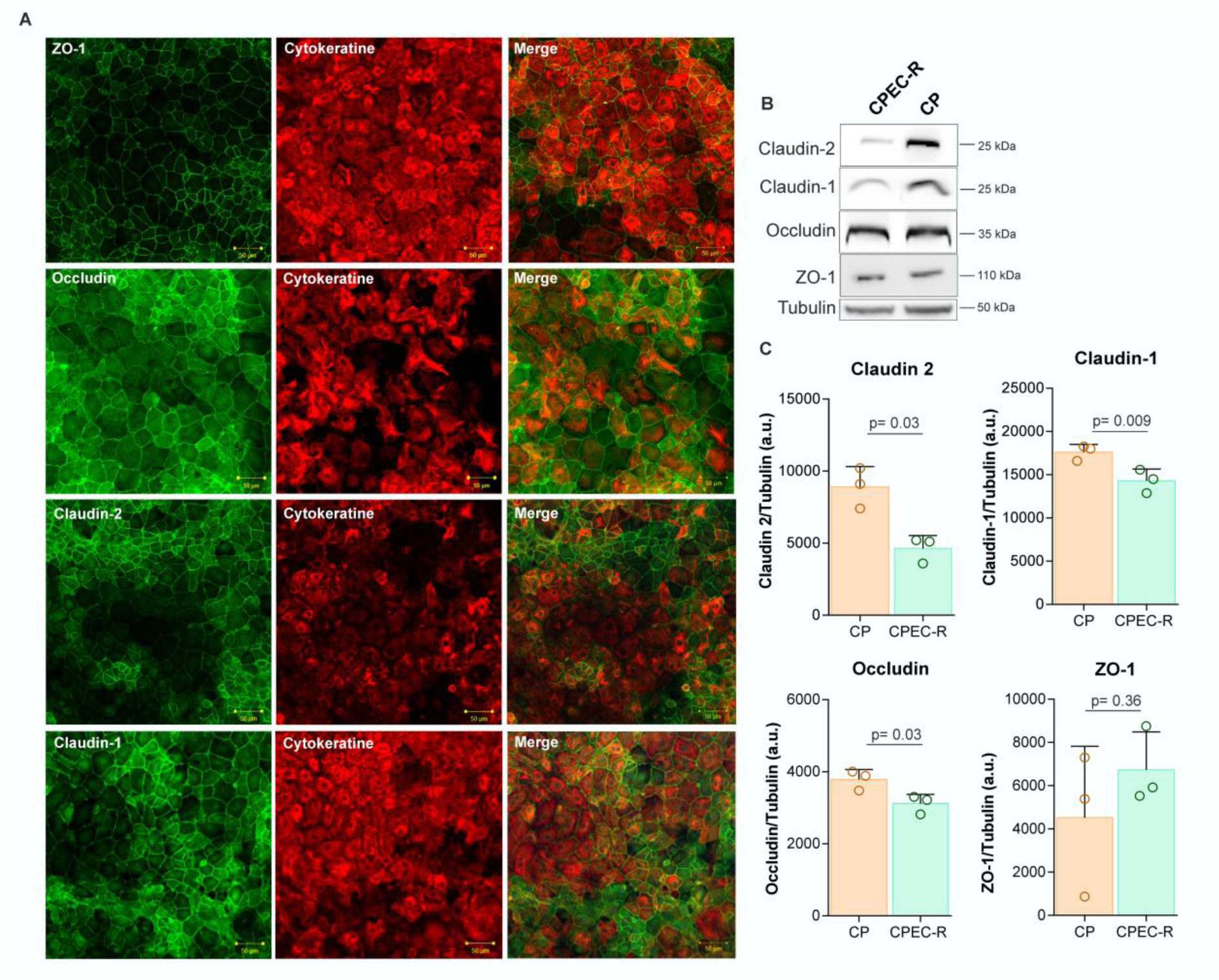
Immunofluorescent and immunoblot analysis of TJ proteins. (**A**) Immunofluorescence images and analysis of TJs for 5-day epithelial monolayer cultivated on Transwell inserts, stained for zonula occludens-1 (ZO-1, green), transmembrane junctional proteins occludin (green), claudin-2 (green) and claudin-1 (green). Colocalization of TJs and cytoplasmic protein pan cytokeratine (red). The staining demonstrated straighter cell junctions and redistribution of cytokeratine. Scale bar 20 µm. (**B**) Western blot analysis of relative concentrations of ZO-1 (110 kDa), occludin (35 kDa), claudin-2 (25 kDa), claudin-1 (25 kDa) and tubulin. (**C**) Bar charts represent relative protein levels of analyzed proteins in choroid plexus compare to *in vitro* epithelial monolayer. Band intensities from three independent experiments (n= 3) were normalised against tubulin values. Values presented are mean ± SD. P < 0.05 was considered to be statistically significant. * p < 0.05, ** p ≤ 0.01 and *** p ≤ 0.001.

### BCSFB barrier expressed pharmaceutically important receptors and influx/efflux transporters

BCSFB is responsible for supplying nutrients, hormones, and drugs between blood and CSF. CP epithelial cells express a wide spectrum of receptors, transporters, and ion channels that play essential roles in the barrier functionality of the BCSFB. Next, we analysed transport-relevant features of barrier, including receptors (LRP-1, LRP-2), influx and efflux drug transport systems, namely ATP-binding cassette (ABC) carrier transporters (BCR-1, MRP1, MRP3, MRP5, Pgp-1, RAGE) and the solute carrier (SLC) group of membrane transport proteins (LAT-1). From a pharmacological point of view, these transporters have a protective role because they limit the entry of neurotoxins into the brain; on the other hand, they hinder effective drug delivery for CNS pharmacotherapy. By immunocytochemistry, we detected the expression of these receptors and transporters in an epithelial monolayer of the inverted BCSFB model (Figure 4A). The transporters are distributed in a polarized fashion at the apical and basolateral surfaces. The polarized distribution enables a regulated, uni- and bi-directional transport of molecules across the BCSFB. Using confocal microscopy, we showed that Pgp-1 was localized mainly in the luminal membrane and Mrp3 in the abluminal membrane (Figure 4B). This demonstrate that CPEC-R cells are polarized in culture. All investigated proteins were also detected on protein level (Figure 4C). By western blot analysis we compared production of all proteins between epithelial monolayer and whole CP tissue (Figure 4D). There was an significant decrease in expression of LRP-2 (CPEC-R: 23964 ± 188.6, CP: 19035 ± 2040, p= 0.012) and MRP3 (CPEC-R: 27919 ± 883.6, CP: 19430 ± 809, p= 0.0021) compare to rat choroid plexus.

**Figure 4.**
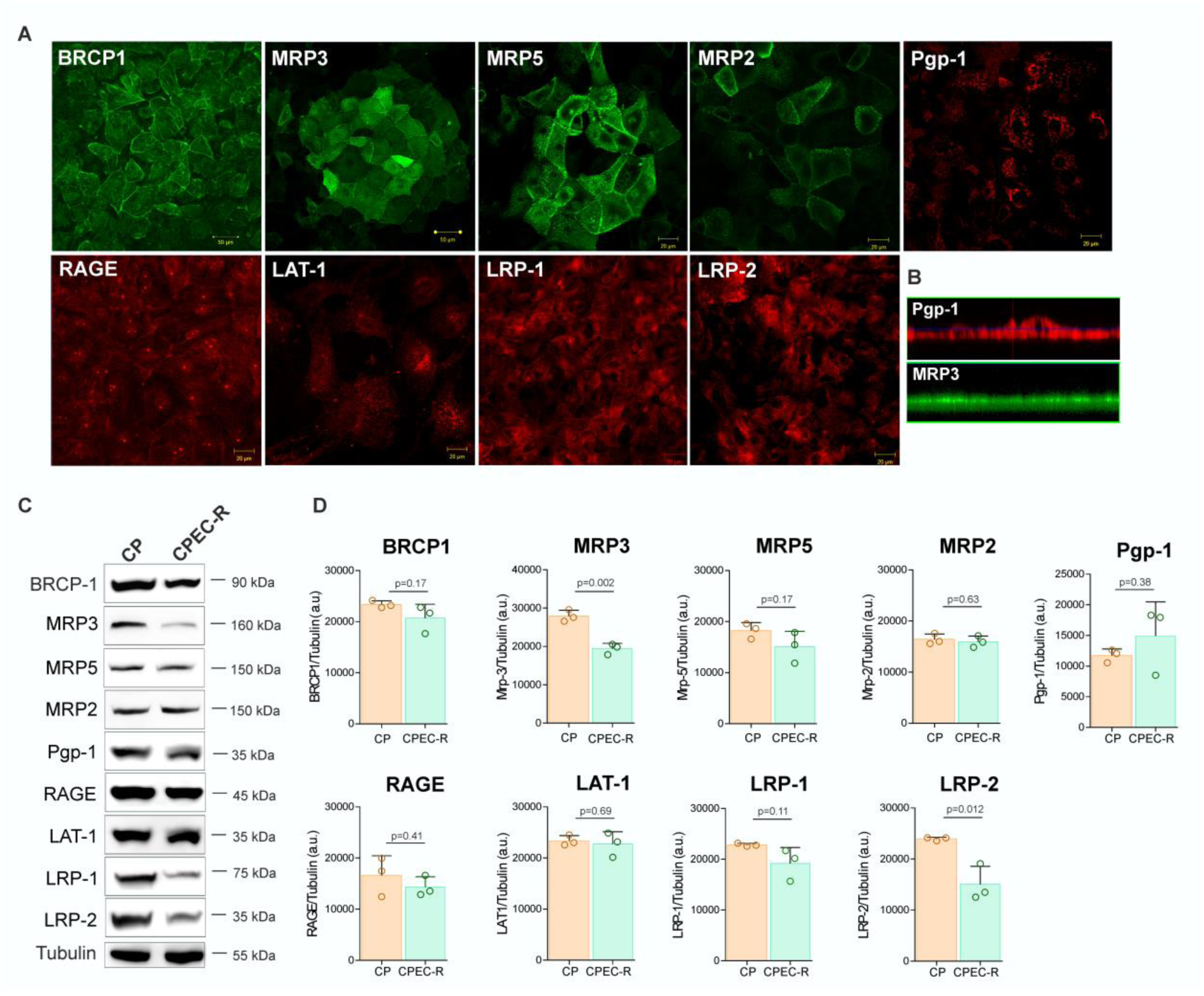
Immunofluorescent and immunoblot analysis transporters. (**A**) Immunofluorescence images of transporters expressed by epithelial monolayer. Scale bar 50 µm. (**B**) Confocal images of epithelial monolayer showing polarized distribution of transporters P-gp1 (shown in red) showed expression within the apical compartment, Mrp1 (shown in green) was expressed in the basolateral membrane. (**C**) Western blot analysis of relative concentrations of analyzed transport proteins. (**D**) Bar charts represent relative protein levels of detected proteins in choroid plexus compared to *in vitro* epithelial monolayer. Band intensities from three independent experiments (n= 3) were normalized against tubulin values. Values presented are mean ± SD. P < 0.05 was considered to be statistically significant. * p < 0.05, ** p ≤ 0.01 and *** p ≤ 0.001.

### CPEC-R actively secrete important proteins

We assessed alterations in protein concentration linked to the production of the secretome by epithelial cells. We analysed secretome samples after 2, 4, 6, and 12 hours and performed statistical testing and over-representation analysis (ORA) (Figure 5A). Complete data are shown in Supplementary file 1. In total we identified 576 proteins. Heatmap reflected differential protein expression in time. Our findings revealed that epithelial cells exhibited low protein production after 2 hours, with a gradual increase in excretion over time. Several proteins showed an increase after 2 hours, however we supposed that these changes were associated with the media change. After a 12-hour period, we observed a significant increase in the amount of secreted proteins including growth factors, cell-matrix proteins, proteases, carrier proteins, inflammation related proteins. In Over-Representation Analysis (ORA), it was anticipated that the categorization of groups according to protein abundance would reveal distinct biological functions. The results of gene ontology analysis according to molecular function (MF) and biological processes (BP) is shown in Figure 5B. Regarding biological processes, proteins involved in extracellular matrix organization, cytoskeleton organization, and collagen fibril organization were identified in the epithelial secretome. Concerning molecular function, we found proteins associated with actin binding, extracellular matrix binding, glycosaminoglycan binding, and heparin binding. These groups of proteins regulate the mechanical properties of tissue as cell proliferation, cell adhesion, growth factor signalling, immune cell function, and collagen structure.

**Figure 5.**
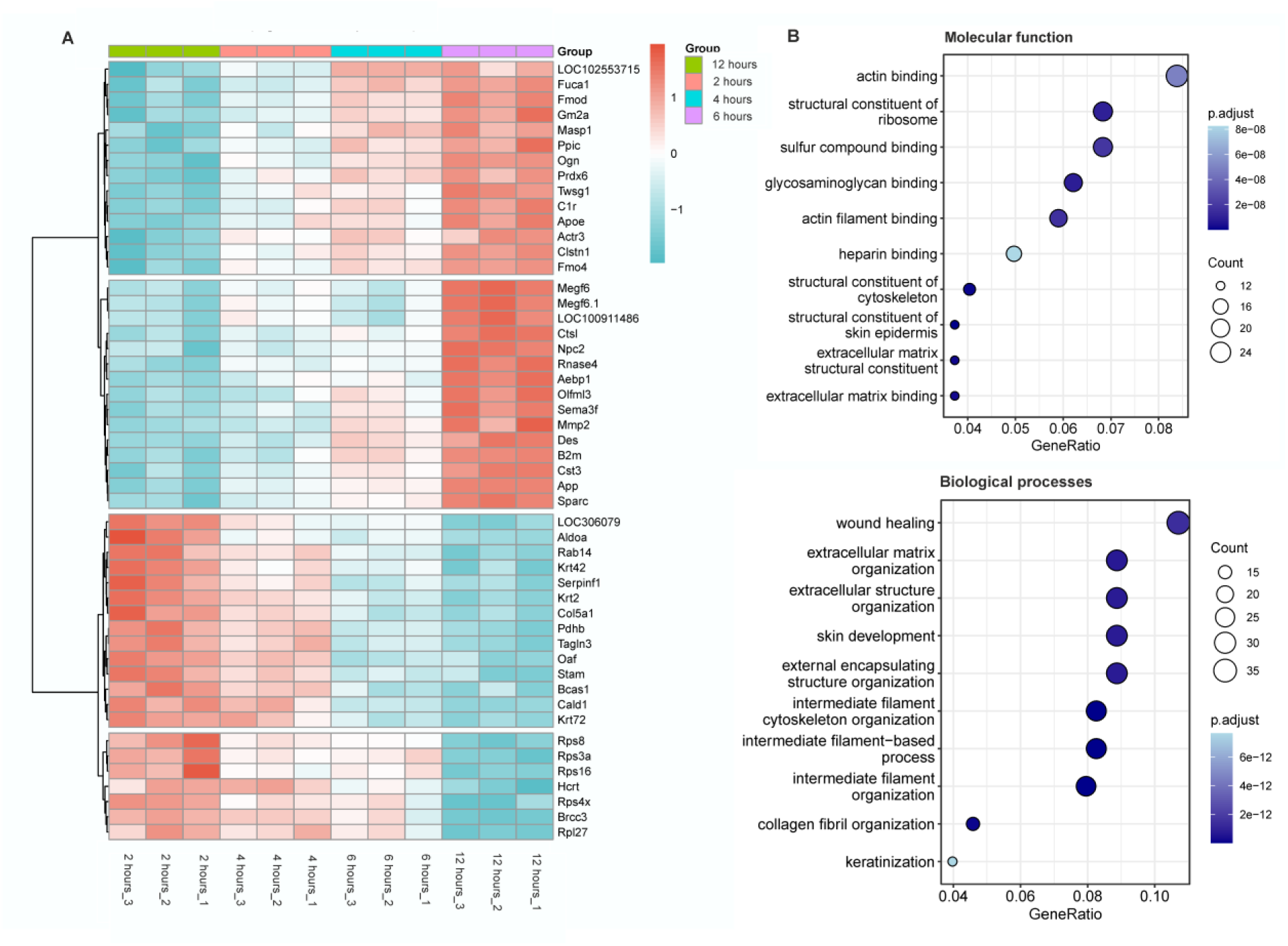
Proteomic analysis of epithelial secretome. (**A**) Hierarchical clustering analysis shown in Heatmap using Ward’s method. Heatmap showing the abundance profile of the proteins secreted into media during the time. (**B**) Gene ontology analysis of the altered proteins. Dotplot, showing the results of ORA, along with the statistical significance (p-value cut off (0.01). Top 10 annotated terms reflecting biological processes (BP) and molecular functions (MF) in the epithelial secretome. The X-axis represents fold enrichment, left Y-axis represents GO terms. p-values ranging from blue (higher significance) to dark blue (lower significance).

## DISCUSSION

Here, we have developed a protocol to establish the primary culture of choroidal epithelial cells (CPEC-R) derived from rats, which were subsequently employed in establishing an inverted model of BCSFB. We showed that isolated CPEC-R displayed a characteristic cuboidal morphology and were positive for choroidal marker, pan cytokeratin. Constitutive expression of cytokeratin showed that the isolated cells adopted epithelial phenotype. To ensure a high yield of epithelial cells, the digestion procedure and sufficient number of cells at the stage of seeding were critical. We found that the well-controlled pronase digest resulted in high viability as well as good cell attachment. The seeding amount higher than 1.10^4^/ml led to healthy growth and fast proliferation. Addition of cytosine arabinoside suppressed the growth of contaminating cells. Finally, epithelial culture developed into a confluent monolayer within 5-6 days.

To design new strategies how to overcome CNS barriers, establish a new delivery system for neuropharmaceuticals, and improve high-throughput drug screening techniques, we developed functionally relevant BCSFB *in vitro* model. Suitable *in vitro* model should mimic *in vivo* environment and demonstrated functional TJs, that are critical for building the selective epithelial barrier. Presence of intercellular TJs (ZO-1, claudin-1, -2 and occludin) resulted to high TEER, restricted paracellular permeability and maintaining the epithelial polarity. Previous BCSFB *in vitro* models recapitulated the characteristic barrier properties, which mainly depend on the formation of TJs (Faich and Stadel 1989, Schroten, Hanisch et al. 2012). Imunofluorescence staining revealed that all investigated TJ were localized abluminaly at the sites of intercellular contact suggesting the formation of highly organised and continuous TJs. Besides that, TJ proteins were detected on protein level. We described a mild decrease of claudin-1 and occludin compare to rat choroid plexus tissue. On the other hand, claudin-2 was dramatically lower expressed in CPEC-R. This indicating a loss of certain characteristics of epithelial cells in primary rat choroidal culture. It has been also shown that expression of claudin-2 in Z310 and TR-CSFB rat epithelial cell lines led to decreased TEER values (Furuse, Furuse et al. 2001). Previously, several rodent epithelial cell lines (Z310, TR-CSFB3) and primary cultures have been analysed regarding the expression of TJ and AJ proteins. Transmission electron microscopy revealed the presence of tight junction (TJ) protein complexes in the sub-apical lateral membranes located between adjacent Z310 and TR-CSFB3 cells.However, immortalized cell lines did not reflect the in vivo situation (Szmydynger-Chodobska, Pascale et al. 2007, Shi, Li et al. 2008, Klas, Wolburg et al. 2010). On the contrary, primary cultures exhibited well-defined junctional complexes characteristic of a mature BCSFB. This was evident through the junctional localization of E-cadherin, as well as claudins-1, -2, -3, and -11 (Klas, Wolburg et al. 2010, Lazarevic and Engelhardt 2016, Lauer, Marz et al. 2019).

To characterize and quantify barrier phenotype we used two standard methods: TEER and paracellular permeability to LY. Presence of intact and continuous TJs agrees with high electrical resistance. The average TEER values were 172 ± 9.5 Ω × cm^2^ and remained at this value for 3 days, indicating the persistent formation of TJs. We also showed that TEER significantly increased after steroid treatment (hydrocortisone, dexamethasone). This is in good correlation with previous results that showed that corticosteroids regulate expression of TJ proteins (Tenenbaum, Matalon et al. 2008). Dexamethasone, functioning as a synthetic glucocorticoid, has demonstrated the ability to enhance barrier strength (Forster, Burek et al. 2008, Stone, England et al. 2019). On the contrary, reducing the serum resulted in a decrease in TEER values. Previously, it was demonstrated that TEER values for primary cultures were significantly higher than those described for cell lines. The decreased TEER values were consistent with the absence of continuous tight junctions. TR-CSFB3 rodent cells reached membrane resistance of 30-50 Ω × cm^2^. For the Z310 rat cell line TEER values varying between 150 - 200 Ω × cm^2^ (Zheng and Zhao 2002). Porcine cell line cultivated on inserts displayed TEER around 600 Ω × cm^2^ (Schroten, Hanisch et al. 2012). Compared to the TEER values observed in our primary rat cells, TEER values in other primary rat cultures were only around 30 Ω × cm^2^. Primary porcine choroid plexus (CP) exhibited comparable values in the range of 100-150 Ω × cm^2^ (Gath, Hakvoort et al. 1997, Baehr, Reichel et al. 2006, Klas, Wolburg et al. 2010). Barrier characteristics of our model were further confirmed by the measurement of permeability for Lucifer yellow (440 Da) as a small molecule tracer. The Pe values for LY in our model were comparable to the Pe values measured for the another impermeable tracers such as inulin, dextran, sucrose, mannitol (Vandenhaute, Stump-Guthier et al. 2015, Drolez, Vandenhaute et al. 2016, Nishihara, Soldati et al. 2020).

The expression of selective transporters was confirmed by immunocytochemistry and western blot. In primary cultures, the expression of ABC and Slc transporters was comparable but lower when compared to intact tissue. We found significantly decreased expression of Mrp3 and LRP-2. In previous study, the transporters were analyzed on mRNA levels. They showed lower expression of ABC transporters (Mrp1, Mrp4, Mrp5) and absent expression of SLC compare to tissue (Choudhuri, Cherrington et al. 2003, Klas, Wolburg et al. 2010).

Taken together in the present study, we have successfully established an inverted *in vitro* model of rat BCSFB with barrier characteristics that closely resemble in vivo situation.. Immunostaining of specific markers and junctional proteins leads to a qualitative confirmation of barrier integrity of an epithelial monolayer. We also showed that this model expresses pharmacologically important transporters and receptors that mediate the selective transport of molecules between blood and CSF. In the future our model can be used for pre-clinical drug discovery and the development of innovative delivery systems for therapeutics or physiological and pathophysiological research studies.

## Acknowledgments

This work was supported by APVV-21-0321 and VEGA 2/0129/21.

## Author contributions

The supervision and design of this study was carried out by A.K. and P.M. Cell isolation and development of BCSFB barrier was done by P.M. and K.H. Western blot and immunocytochemistry was done by P.M. J.V. provide animals for experiments. Visualization and interpretation of the results were done by P.M, A.K., and K.H. The final version of article was edited and reviewed by all co-authors.

## Ethical approval

All animal experiments were performed according to the institutional animal care guidelines and in accordance with international standards (Animal Research: Reporting of In Vivo Experiments guidelines) and approved by the State Veterinary and Food Administration of the Slovak Republic (Ro-933/19-221/3a) and by the Ethics Committee of the Institute of Neuroimmunology, Slovak Academy of Sciences.

## Competing interests

The authors claim no competing interests.

## Data availability

Other questions can be directed to the corresponding authors.

## Notes

### Competing Interest Statement

The authors have declared no competing interest.

